# Evidence from the field that multiple processes maintain hidden adaptive capacity to a novel environment in a wild daisy

**DOI:** 10.64898/2026.08.01.741386

**Authors:** Greg M. Walter, Delia Terranova, Giuseppe Emma, James Clark, Salvatore Cozzolino, Simon J. Hiscock, Antonia Cristaudo, Jon Bridle

**Affiliations:** University of Bristol, School of Biological Sciences, Bristol BS8 1TQ, UK; Monash University, School of Biological Sciences, Melbourne 3800, Australia; University of Tasmania, School of Natural Sciences, Hobart, 7005, Australia; University of Naples Federico II, Department of Biology, Naples 80126, Italy; University of Catania, Department of Biological, Geological and Environmental Sciences, Catania 95128, Italy; University of Oxford, Department of Plant Sciences, Oxford, OX1 3RB, UK; University of Bath, Milner Centre for Evolution, Bath, BA2 7AY, UK; University College London, Department of Genetics, Evolution and Environment, London WC1E 6BT, UK

**Keywords:** adaptive potential, balancing selection, recessive mutations, seasonal variation, cryptic genetic variation

## Abstract

Populations often persist in novel environments despite predictions that adaptive capacity to such conditions should be limited by a lack of genetic variation. A leading hypothesis is that genetic variation for adapting to novel environments is maintained but remains hidden (or ‘cryptic’) under native conditions. However, direct evidence of mechanisms that maintain this adaptive potential in populations is scarce. We integrate data from four large-scale field experiments to test whether variation in selection across environments and life history, together with genetic architecture, maintains genetic variation important for adapting to novel environments. Using a quantitative genetic breeding design, we generated families of the Sicilian daisy, *Senecio chrysanthemifolius* (Asteraceae), and planted seeds and cuttings across native and novel elevations on Mount Etna. We tracked fitness across elevations, life stages, seasons and generations. Genotypes with higher survival and flowering success at the novel elevation increased adaptive potential, but were not strongly disadvantaged in the native environment. A negative genetic correlation in seedling survival across seasons indicated that different genotypes were favoured across temporal variation in native environments. Crosses between genotypes with low and high fitness in the novel environment revealed that genotypes that increased adaptive potential had heritable effects on plasticity and fitness across generations, but were partially recessive and likely to hide in heterozygotes. Together, these results provide rare field-based evidence that inefficient selection in native environments and dominance maintain cryptic adaptive potential for novel environments, while temporal variation in selection maintains genetic variation in fitness more broadly.

## Introduction

From warming climates to novel habitats created by human activity, populations worldwide are encountering environmental change fast enough to threaten their persistence (Parmesan and Yohe 2003; Nadeau et al. 2017; Edelsparre et al. 2024). Whether populations survive such change depends on how quickly they can adapt, which will be determined by the amount of genetic variation in fitness available for natural selection to act on (Gomulkiewicz and Holt 1995; Lande and Shannon 1996). Theory predicts that adaptation to novel environments should be limited by the low availability of genetic variation that is beneficial in the new environment (Bürger and Lynch 1995), yet many populations show a surprising capacity to persist, and in some cases adapt, to novel environments (McNeilly 1968; Antonovics and Bradshaw 1970; Bradshaw 1991; Bell and Gonzalez 2009; Bell 2017; Kreiner et al. 2019; Balstad et al. 2026). This mismatch between theoretical expectation and empirical observations raises a fundamental question: how is genetic variation that enables adaptation to novel environments maintained in natural populations? While research has focused on the mechanisms that maintain genetic variation in populations in general (Charlesworth 2015; Bernatchez 2016; Sharp and Agrawal 2018), less attention has been paid to genetic variation that increases adaptive potential in novel or changing environments, despite such information being critical for predicting the persistence of populations under rapid global change (Dobzhansky 1955; Gomulkiewicz and Holt 1995; Urban et al. 2024; Chevin and Bridle 2025).

During adaptation, natural selection increases the frequency of advantageous alleles while reducing the frequency of alternative alleles, thereby reducing genetic variation. Balancing selection counteracts this loss of genetic variation when different alleles are favoured under different conditions (e.g., across seasons or life stages), which is a leading explanation for how genetic variance in fitness is maintained in populations (Dobzhansky 1955; Gillespie and Turelli 1989; Ruzicka et al. 2026). Across a range of taxa, studies have shown that genetic variation can be maintained by variation in selection across life-history stages (Christie et al. 2018), fluctuating selection across time and space (Anderson et al. 2013; Bergland et al. 2014; Abdul-Rahman et al. 2021), and environment-dependent genetic effects (i.e., genotype-by-environment interactions) on fitness (Carley et al. 2021). In addition, aspects of genetic architecture, such as recessive alleles that are masked in heterozygotes, can reduce the efficiency of selection and allow genetic variation to persist within populations (Charlesworth 2015). However, these mechanisms are typically studied in isolation or under laboratory conditions, and so it remains unclear whether they act together to maintain genetic variation that underlies adaptive potential for novel environments in the field.

Persistence in novel environments may depend on standing genetic variation whose fitness effects are not expressed under native conditions but become advantageous when environments change (Masel 2006; Le Rouzic and Carlborg 2008; Hayden et al. 2011; Ledón-Rettig et al. 2014). In such situations, genotypes that show less severe declines in fitness in novel environments will help to maintain population size, allowing the population to persist and then adapt to the novel conditions (Lande 1988; Lande and Shannon 1996; Bell 2013). Such ‘cryptic’ genetic variation is hidden in native environments and only emerges in novel conditions, resulting in genotypes varying in their response to novel environments that increases genetic variance in fitness and therefore adaptive potential (Chevin et al. 2010; Gomulkiewicz and Shaw 2013; Shaw and Shaw 2014). Exposing genotypes to novel environments is required to reveal cryptic genetic variation, which then allows exploration into how such genetic variation is maintained in the population under natural conditions (Gibson and Dworkin 2004; Paaby and Rockman 2014). Although there is growing empirical support for increased genetic variance in novel environments (Parsons 1987; Sgrò and Hoffmann 1998; Torres-Martínez et al. 2019; Walter et al. 2023; Walter et al. 2026b), the processes that maintain the standing genetic variation underlying such adaptive potential remain poorly understood.

The availability of adaptive genetic variation in novel environments will depend on how selection varies across life-history stages or across time and space, as well as on the underlying genetic architecture of fitness (Dobzhansky 1955; Ruzicka et al. 2026). If alleles that are beneficial in novel environments are not strongly selected against in native environments, selection in native environments may be inefficient at removing genetic variation that contributes to adaptation under novel conditions. This could occur when selection is strongest at early life-history stages and a weak genetic coupling across life-history stages limits the ability for selection to remove genotypes that reduce fitness at later life stages (weak selection hypothesis; Stearns 1989; Roff and Fairbairn 2007; Anderson et al. 2013). Similarly, temporal or spatial environmental heterogeneity can generate variation in selection that favours different genotypes in different ecological contexts (e.g., across seasons or environments), preventing the loss of alleles that are intermittently deleterious in the native range, but that could be beneficial in a novel environment (fluctuating selection hypothesis; Gillespie 1974; Charlesworth et al. 1997; Bürger and Gimelfarb 2002; Culumber et al. 2015; Wittmann et al. 2017; Abdul-Rahman et al. 2021). Genetic architecture can also maintain adaptive variation if alleles with beneficial effects in novel environments are recessive and remain hidden in heterozygotes in native environments (recessive alleles hypothesis; Haldane 1937; Orr and Betancourt 2001; Charlesworth 2015). While each mechanism has been demonstrated in a variety of systems, they have rarely been examined together and are seldom linked to the maintenance of genotypes that increase fitness in novel environments. We therefore lack direct empirical evidence to assess how adaptive potential is maintained in natural populations (Anderson et al. 2014; Kulbaba et al. 2019; Shaw 2019; Peschel et al. 2020).

Testing the mechanisms that maintain adaptive potential requires assaying large numbers of genotypes to estimate additive genetic variance in fitness across multiple environments and life stages, which is logistically challenging. For this reason, most studies measure adaptive potential at a single life stage, in a single environment, or within a single generation, limiting our ability to test how multiple mechanisms contribute to maintaining adaptive capacity. We use *Senecio chrysanthemifolius* (Asteraceae), a self-incompatible, short-lived perennial daisy native to c.400-1500m elevation on Mount Etna (Sicily, Italy) that relies on generalist insect pollinators (e.g., hoverflies). A closely related species, *S. aethnensis,* is endemic to old lava flows at higher elevations (Brennan et al. 2009; Walter et al. 2020). The two *Senecio* species show adaptation to their contrasting habitats associated with differences in plasticity, and in genetic variance in leaf traits (Walter et al. 2022; Walter et al. 2024). In previous work, we found that *S. chrysanthemifolius* showed increased adaptive potential at a novel 2000m elevation for both seedling survival and flowering success (Walter et al. 2023; Walter et al. 2026b), but why this genetic variation persists in the population under native conditions remains unclear. Here, we integrate these previously published fitness measurements within a multivariate framework to test whether selection is inefficient at removing these genotypes from the population. We then include new data from experiments in a different season (Autumn), and across generations, to test whether fluctuating selection across seasons or dominance help to maintain genetic variation important for increasing adaptive potential in a novel environment.

In 2018, we generated 104 families of *S. chrysanthemifolius,* which we transplanted as cuttings (2018, experiment 1) and as seeds (2019, experiment 2). **Weak selection hypothesis:** connecting fitness estimates from experiments 1 and 2, we predicted that additive genetic correlations linking performance across both life stages (seedling survival and flowering success) and elevations (native and novel) would be zero, or weak and negative (rather than strong and negative), which would limit the efficiency of selection at the native site in removing genotypes that increase adaptive potential in the novel environment. To test this, we took a multivariate approach using the full matrix of pairwise genetic correlations among all four fitness measures (survival and flowering success at both elevations), rather than relying on any single correlation in isolation. **Fluctuating selection hypothesis:** planting the same families as seeds in Spring and Autumn (2019, experiment 3) at the native site, we predicted a negative additive genetic correlation in seedling performance across seasons, indicating that different seasons favour different genotypes, with Autumn performance potentially predicting adaptive potential at the novel high elevation. **Recessive alleles hypothesis**: we compared genotypes identified in experiment 1 that showed consistently higher relative fitness in the novel environment (AP, ‘Adaptive Potential’ genotypes) to those with higher relative fitness in the native environment (HR, ‘Home Range’ genotypes) (Walter et al. 2026a). In experiment 4 (2020), we made crosses within (AP×AP and HR×HR) and between (AP×HR) AP and HR genotypes. We grew offspring in the common garden and transplanted cuttings of multiple genotypes per cross-type at each of four elevations on Mount Etna. Crossing *within* AP and HR genotypes tests whether the fitness and phenotype benefits of AP genotypes at a novel elevation are heritable. Crosses *between* AP and HR genotypes then test whether ‘hybrid’ AP×HR genotypes show similar patterns of fitness and phenotype to the HR genotypes, suggesting that AP genotypes remain hidden as recessive alleles within the native range.

## Results

### Weak selection hypothesis

*Genetic variation for adaptive potential is not strongly disadvantaged in the native environment*

We estimated genetic correlations to link genetic variation in performance across elevations and life-history traits. As expected for a quantitative genetic design in the field, individual pairwise correlations were estimated with substantial uncertainty, and all posterior distributions are presented in **Table 1A**. Survival and flowering success showed no clear genetic correlation at the native elevation (r_g_ = -0.12, P(r_g_<0) = 0.66; **Fig. 1A**) but were more genetically integrated at the novel elevation (r_g_ = 0.35, P(r_g_>0) = 0.92; **Fig. 1B**), suggesting that survival and flowering success become more genetically coupled once genotypes are exposed to novel conditions. Genetic correlations linking life stages across elevations showed opposite signs: survival at the native elevation tended to predict higher flowering success at the novel elevation (r_g_ = 0.28, P(r_g_>0) = 0.84; **Fig. 1C**), whereas flowering success at the native elevation tended to predict lower survival at the novel elevation (r_g_ = -0.28, P(r_g_<0) = 0.85; **Fig. 1D**).

**Fig. 1.**
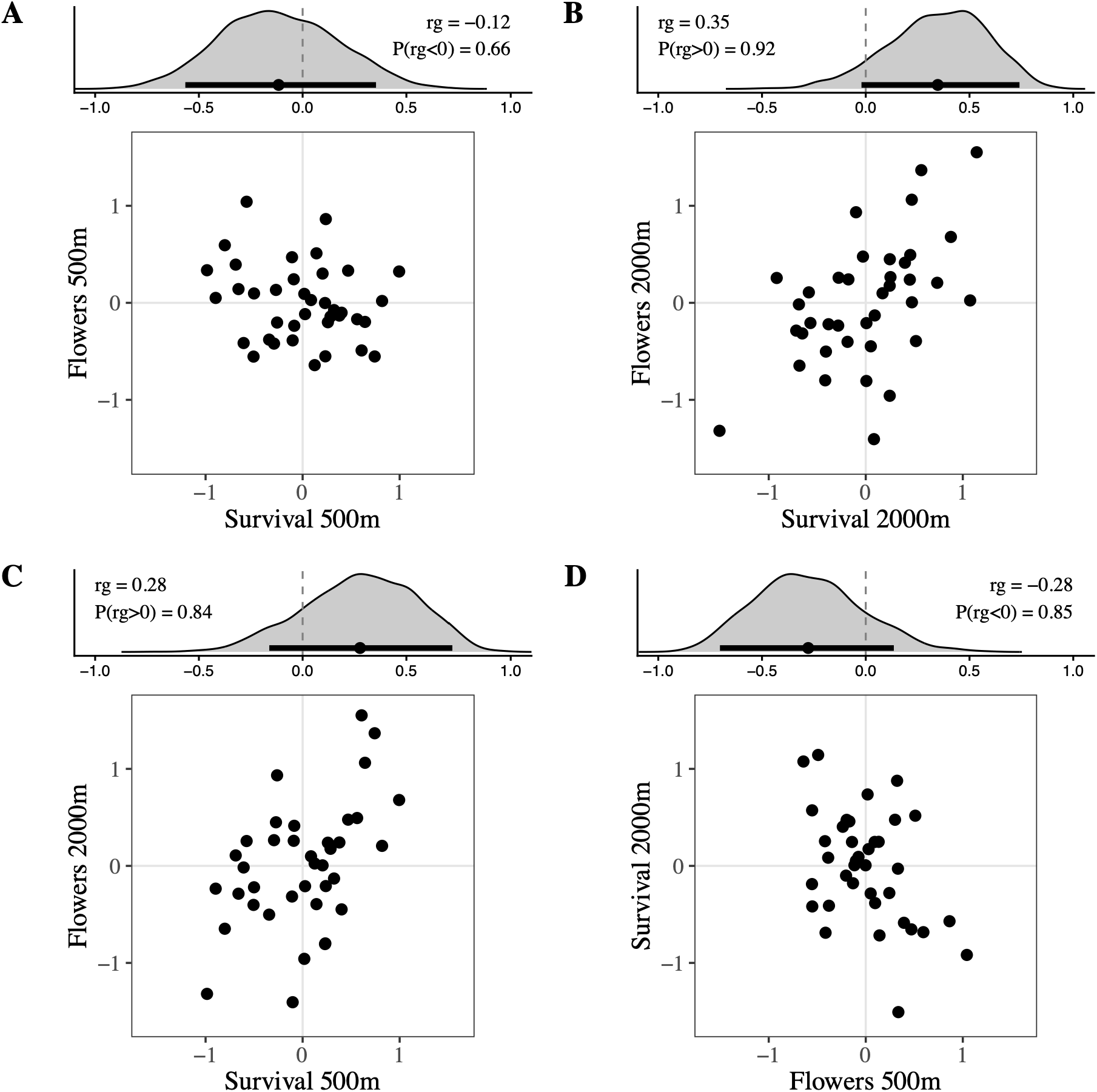
Presenting the additive genetic correlations across life-history traits at each elevation (**A-B**), and across life-history traits and elevations (**C-D**) from **Table 1**. Plots show posterior distributions of the genetic correlation (upper panels: grey frequency density plot with the black bar showing the 90% HPD interval), and sire breeding values (BLUPs, the Best Linear Unbiased Prediction) for fitness (lower panels) underlying the genetic correlations. Survival and flowering success showed no clear genetic correlation at the native 500m (**A**), but were positively correlated at the novel 2000m (**B**). Across elevations, seedling survival at 500m tended to be positively correlated with flowering success at 2000m (**C**), whereas flowering success at 500m tended to be negatively correlated with seedling survival at 2000m (**D**). **Fig. S2** shows the 90% HPD intervals for each sire BLUP, illustrating fitness differences among sires.

**Table 1.** (A) Additive genetic correlation matrix in field fitness showing the genetic correlations between seedling survival and flowering success between the native (500m) and novel (2000m) elevations. Presented are the posterior means with 90% Highest Posterior Density (HPD) intervals in brackets below the diagonal. Above the diagonal are the probabilities of direction (pd), i.e., P(rg<0) or P(rg>0) depending on the sign of the correlation. **(B)** The loadings of the first axis of the genetic correlation matrix show the contribution of each of the performance (across life-history traits and elevations) measures to describing the major axis (***g***max, λ=1.84, 1.17–2.50 90% HPD, 46.1% of total variance explained) of variation in fitness.

| (A) |  |  |  |  | (B) |
| --- | --- | --- | --- | --- | --- |
| | Flowers 500m | Flowers 2000m | Survival 500m | Survival 2000m | $g_{\text{max}}$ |
| Flowers 500m | — | 0.68 | 0.66 | 0.85 | 0.33 |
| Flowers 2000m | -0.10 [-0.49, 0.27] | — | 0.84 | 0.92 | -0.47 |
| Survival 500m | -0.12 [-0.57, 0.35] | 0.28 [-0.16, 0.72] | — | 0.95 | -0.54 |
| Survival 2000m | -0.28 [-0.70, 0.14] | 0.35 [-0.02, 0.74] | 0.48 [0.08, 0.87] | — | -0.61 |

To test whether patterns of genetic correlations reflect shared genetic variation, we took a multivariate approach to quantify how genetic variation in fitness was shared across elevations and life-history traits. By decomposing the genetic correlation matrix (**Table 1A**) into orthogonal axes (analogous to principal components analysis), we identified the dimensions along which genetic variation in fitness was most correlated. Using a permutation test to create a suitable null distribution, we found statistical support for the leading axis, ***g***_max_ (**Fig. S1**), which captured the greatest amount of shared genetic variation in fitness (λ=1.84, 46.1% of total variance). Flowering success at the native elevation loaded in one direction, while survival at the native elevation along with survival and flowering at the novel elevation loaded in the opposite direction (**Table 1B**). This suggests that there was little shared additive genetic variation in fitness between native flowering and the other three performance measures.

This lack of shared genetic variation between native flowering and the other three performance measures is also apparent when comparing correlations directly (**Table S1**). All three correlations involving native flowering were weak, tended to be negative, and each was lower than the three positive correlations in **Table 1** (nine non-independent comparisons; P(Δ<0) = 0.83–0.97, where Δ is the difference between the correlations; **Table S1**). The clearest difference involved the correlation between native flowering and novel survival (r_g_ = -0.28), which was credibly lower than correlations between survival across elevations (r_g_ = 0.48; P(Δ<0) = 0.97), between survival and flowering at the novel elevation (r_g_ = 0.35; P(Δ<0) = 0.96), and between native survival and novel flowering (r_g_ = 0.28; P(Δ<0) = 0.92) (**Table S1**). The weak correlation between native and novel flowering (r_g_ = −0.10) was also lower than the correlation between survival across elevations (r_g_ = 0.48; P(Δ<0) = 0.94). This indicates detectable structure in the correlation matrix, consistent with the axis identified by the permutation test above.

Together, these results suggest that genetic variation for flowering success at the native elevation is decoupled from genetic variation underlying survival at both elevations, and flowering at the novel elevation. Consistent with the weak selection hypothesis, we find no evidence that selection at the native site is strong enough to remove genotypes that increase adaptive potential at the novel elevation. Although our data cannot distinguish whether this reflects weak antagonistic selection at the native site or the absence of selection altogether, both scenarios are consistent with these genotypes persisting in the native population.

### Fluctuating selection hypothesis

*Seasonal selection favoured different genotypes but was not clearly linked to performance in the novel environment*

We found a moderately strong negative genetic correlation between seedling survival in Spring versus Autumn (r_g_ = -0.49; P(r_g_<0) = 0.92; **Fig. 2A**), which suggests that different genotypes showed higher survival across seasons. However, we found no clear evidence that Autumn seedling survival predicted flowering success (r_g_ = -0.18, P(r_g_<0) = 0.69; **Fig. 2B**) or seedling survival (r_g_ = 0.16, P(r_g_>0) = 0.65; **Fig. 2C**) at 2000m. Both correlations were weak and uncertain, indicating that genotypes with better survival in Autumn were not necessarily the highest performing genotypes at 2000m.

**Fig. 2.**
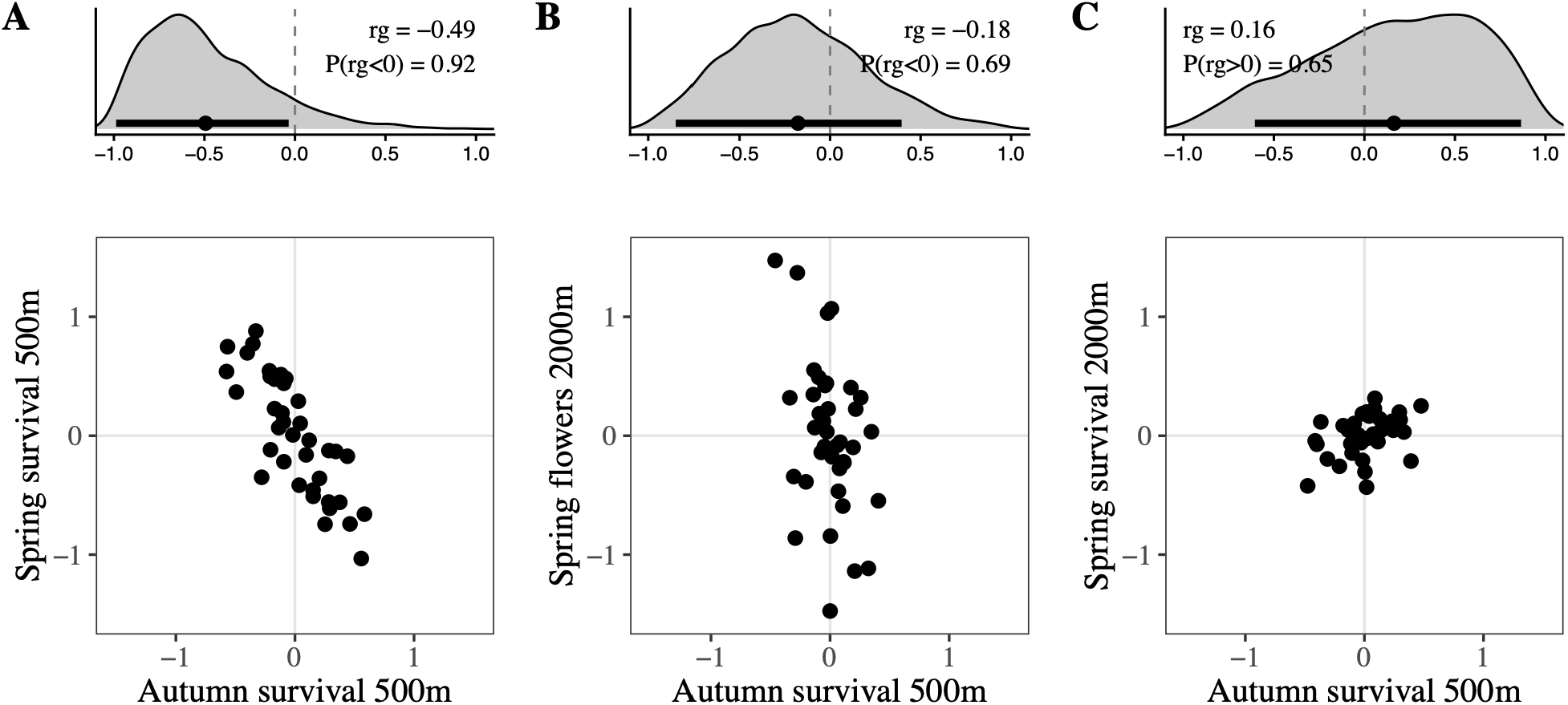
Genetic correlations across seasons with the posterior distribution presented in the upper panels (grey frequency density plot with the black bar showing 90% HPD), and the associated sire breeding values in the lower panels. **(A)** A negative genetic correlation in seedling survival between Autumn and Spring at the native elevation suggests different alleles were favoured across seasons. However, Autumn seedling survival showed no clear evidence of predicting fitness at 2000m for either **(B)** flowering success, or **(C)** seedling survival.

### Recessive alleles hypothesis

*Genotypes with high fitness in novel environments are partially recessive and retain fitness and phenotype benefits across generations*

*Fitness:* Crossing within genotype classes showed that genotypes with higher relative fitness at the novel 2000m elevation (AP, ’Adaptive Potential’ genotypes) and those with higher relative fitness at the native elevation (HR, ’Home Range’ genotypes) produced offspring with fitness patterns similar to their parents, indicating heritable differences in fitness between AP and HR genotypes (**Fig. 3**).

**Fig. 3.**
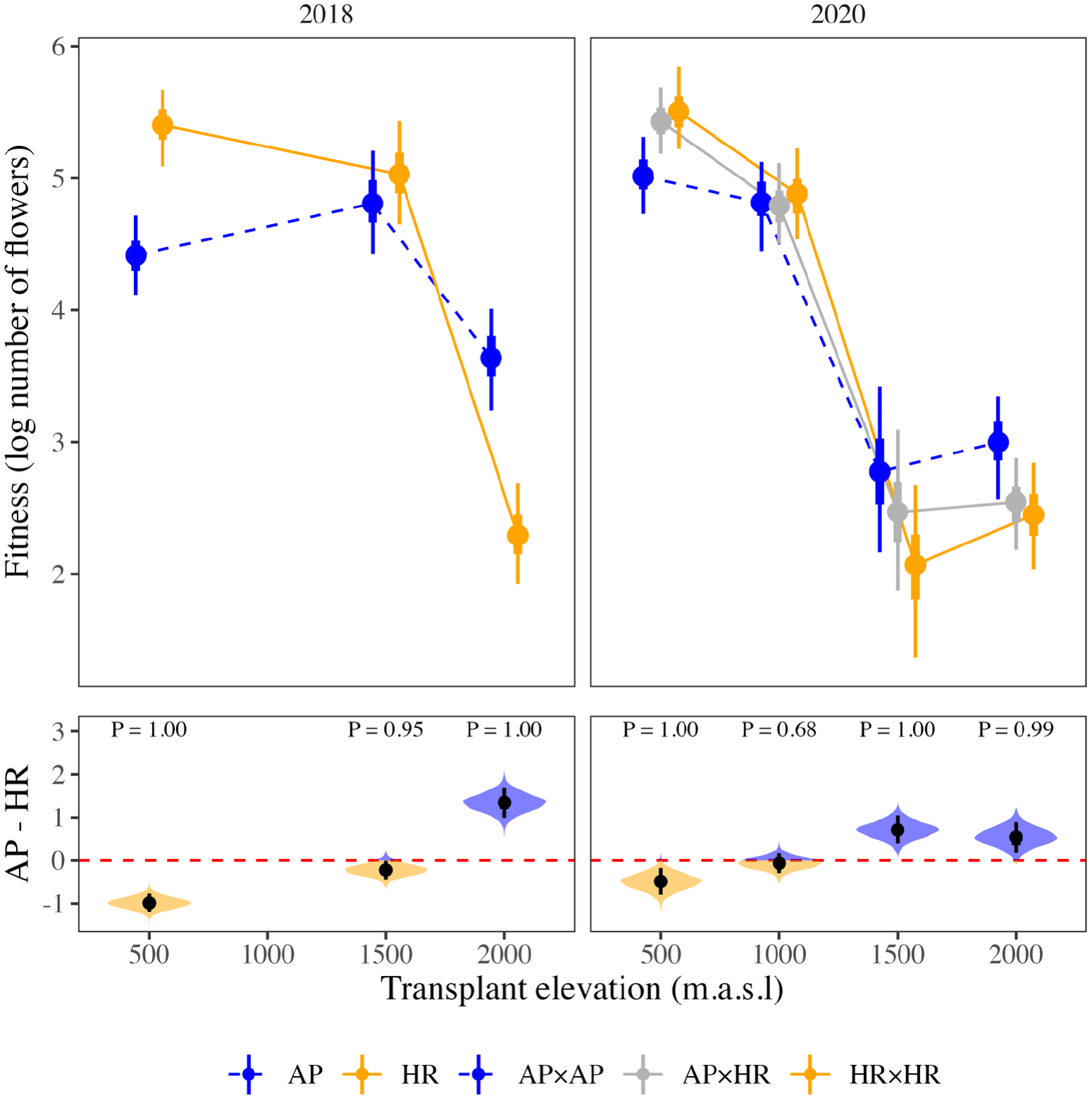
Fitness of AP and HR genotypes in 2018, and of their offspring when crossed within and between genotype classes and transplanted as cuttings in 2020. Upper panels: points represent the posterior median fitness (log number of flowers) for each genotype class at each elevation, with error bars showing the 50% (thick) and 90% (thin) HPD credible intervals. Lower panels: posterior distribution of the difference in fitness between AP and HR genotypes (2018), and between AP×AP and HR×HR genotypes (2020), at each elevation. Black points and bars show the median with 50% and 90% HPD intervals, and the red dashed line marks zero (no difference). Shading shows where AP (blue) or HR (orange) genotypes had higher fitness, and P is the probability of direction (pd), i.e., P(Δ>0) or P(Δ<0) depending on the sign of the difference. AP genotypes had lower fitness than HR genotypes at the native 500m elevation but higher fitness at the novel 2000m elevation, resulting in credible Cross×Elevation interactions between the native and novel elevations in both years (**Table S2**).

This was supported by credible Cross×Elevation interactions between the native and novel elevations (i.e., the AP–HR difference at 500m minus that at 2000m) in both 2018 (-2.33, -2.71–-1.93 90% HPD; P(<0) = 1.00) and 2020 (-1.04, -1.50–-0.59 90% HPD; P(<0) = 1.00; **Fig. 3**; **Table S2**). Furthermore, most pairwise interactions across elevations were credible (**Table S2**), suggesting broad differences in the fitness response of AP and HR genotypes to elevation.

AP×HR hybrid genotypes showed similar patterns of fitness to HR×HR genotypes at both native and novel elevations (**Fig. 3**). To test whether this reflects dominant genetic effects of HR over AP alleles, we calculated the potence ratio (Falconer 1960; de Vienne and Fievet 2020), a dominance index that quantifies the deviation of AP×HR from the midparent value: zero represents purely additive effects, while values of 1 represent complete dominance towards HR×HR. At both elevations, AP×HR deviated from the midparent value towards HR×HR with a potence ratio of 0.70 (-0.44–2.17 90% HPD; P(>0) = 0.90) at the native site, and 0.66 (-0.43–2.28 90% HPD; P(>0) = 0.89) at the novel elevation. These results therefore suggest partial dominance of HR over AP. Both directions of the cross (i.e., AP×HR and HR×AP) showed similar fitness at all elevations (**Fig. S3**), suggesting no effect of directionality on fitness.

*Leaf traits:* AP×AP genotypes tended to have smaller, less complex leaves with more leaf indents and higher Specific Leaf Area (SLA) than HR×HR genotypes, with differences credible at most elevations (**Fig. 4A**), suggesting strong phenotypic differences between AP and HR genotypes. We found credible Cross×Elevation terms for leaf area, complexity and SLA between the native site and novel elevation, indicating that AP×AP and HR×HR crosses differed in plasticity (i.e., their sensitivity to the environment) across elevations (Cross×Elevation for leaf area: -0.25, -0.44**–**-0.05 90% HPD, P(<0) = 0.98; leaf complexity: -0.10, -0.18**–**0.00, P(<0) = 0.95; SLA: 0.05, 0.00**–**0.12, P(>0) = 0.94; **Fig. 4A; Table S3**). For leaf area and complexity, AP×AP genotypes showed lower plasticity than HR×HR crosses at the two higher elevations (**Fig. 4B**), which supports the original study where higher fitness at the 2000m novel elevation was associated with lower plasticity (Walter et al. 2023). By contrast, AP×AP genotypes showed a stronger decrease in SLA than HR×HR genotypes at the novel elevation (**Fig. 4B**). Many of the pairwise interactions across elevations were credible (**Table S3**), suggesting broad differences between AP and HR genotypes in their plastic responses to elevation.

**Fig. 4.**
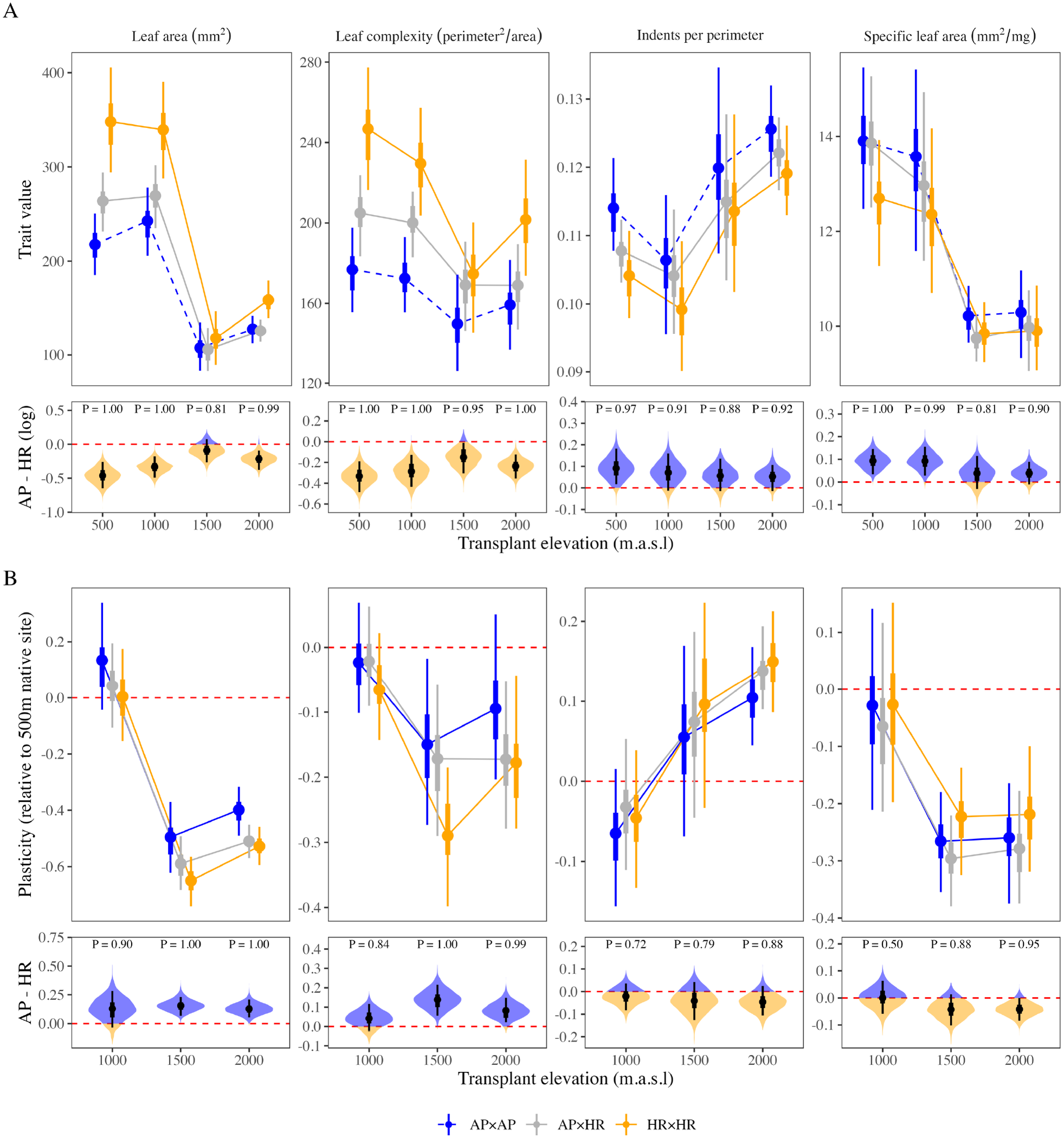
The phenotypic effects of crossing within and between AP and HR genotypes. **(A)** Leaf traits for each cross type at each elevation. Points represent the posterior median, with error bars showing the 50% (thick) and 90% (thin) HPD credible intervals. Lower panels show the posterior distribution of the difference between AP×AP and HR×HR genotypes at each elevation on the log scale (i.e., the log ratio of their trait values). The interaction between cross (AP×AP and HR×HR) and elevation (500m–2000m) was credibly different from zero for leaf area, complexity and SLA (**Table S3**). AP×HR genotypes were often phenotypically intermediate between AP×AP and HR×HR. **(B)** Plasticity of each genotype, calculated as the change in trait value from the native 500m site to each elevation, relative to the native site. Lower panels show the posterior distribution of the difference in plasticity between AP×AP and HR×HR genotypes. Leaf area and complexity were the only traits that showed consistent credible differences in plasticity between AP×AP and HR×HR genotypes, and AP×HR crosses tended to show similar plasticity to HR×HR crosses. In all lower panels, black points and bars show the median with 50% and 90% HPD intervals, the red dashed line marks zero (no difference), shading shows where AP×AP (blue) or HR×HR (orange) genotypes had the higher value, and P is the probability of direction (pd), i.e., P(Δ>0) or P(Δ<0). **Fig. S5** presents the estimates for each individual genotype, showing variation within each cross type.

AP×HR hybrid genotypes tended to show intermediate phenotypes to AP×AP and HR×HR (**Fig. 4A**). At the novel elevation, however, AP×HR genotypes showed similar trait values to AP×AP genotypes for leaf area and complexity (**Fig. 4A**), which is the opposite pattern to what we found for fitness. By contrast, AP×HR genotypes showed similar patterns of plasticity to HR×HR crosses at the novel elevation (**Fig. 4B**), supported by credible potence ratios of 0.73 (-0.24–1.78 90% HPD; P(>0) = 0.94) for leaf area, and 0.86 (-0.63–2.67 90% HPD; P(>0) = 0.91) for leaf complexity, which indicates dominance towards HR despite substantial uncertainty. This suggests that alleles underlying AP genotypes had partially recessive genetic effects on plasticity that are consistent with the patterns found for fitness (**Fig. 3**), despite showing largely intermediate trait values (**Fig. 4A**).

## Discussion

We previously showed that adaptive potential increases in a novel compared to a native environment at early and later life-history stages (Walter et al. 2023; Walter et al. 2026b). In this study, we used field measures of fitness across life stages, environments and generations to test mechanisms that could maintain standing genetic variation in a population that increases adaptive potential in novel environments. Our results provide support for all three hypotheses, and suggest ways that selection in the native environment is unlikely to remove genotypes that increase adaptive potential for novel environments. Consistent with the **weak selection hypothesis**, genotypes with higher fitness in the novel environment showed higher seedling survival at both elevations, but were not strongly disadvantaged for flowering success at the native elevation (**Table 1**, **Fig. 1**). This suggests that selection on flowering success in the native environment is unlikely to remove genotypes that contribute to adaptive potential in the novel environment. In support of the **fluctuating selection hypothesis**, seasonal variation in selection at the native elevation favoured different genotypes (**Fig. 2A**), but we found no clear evidence that performance in Autumn predicted performance at the novel high elevation (**Fig. 2B-C**). Fluctuating selection may therefore maintain adaptive potential more broadly, but without a simple or direct link between genotypes selected across seasons and fitness in novel environments. By crossing within versus among genotypes that showed differences in fitness at the novel elevation, we found support for the **recessive alleles hypothesis**. Differences between AP and HR genotypes were heritable, and AP genotypes showed partially recessive effects on both fitness (**Fig. 3**) and plasticity (**Fig. 4B**), but were intermediate for trait means (**Fig. 4A**). This suggests that genetic variation in plasticity underlies adaptive potential at the novel elevation, which is masked in heterozygotes in the native range, providing empirical support for the role of partially recessive alleles in maintaining adaptive potential.

Overall, our field data indicate that weak selection and partial recessivity allow genotypes favoured in the novel environment to persist in the population, while fluctuating selection across seasons maintains adaptive potential more broadly. Although we show that relative performance differed across life-history stages and elevations, we cannot easily infer the strength of selection on specific variants, or how underlying genetic architecture (e.g., pleiotropy) determines the number of loci contributing to the fitness patterns we observed. This is a prime avenue for future research.

### Weak selection and partial recessivity shelter genetic variation for adaptive potential

When populations are close to their fitness optima, selection should be weak and inefficient at removing alleles with small or context-dependent effects, increasing the scope for mechanisms of balancing selection to maintain genetic variation (Turelli 1984; Barton and Keightley 2002; Charlesworth 2015). Our results support this idea: by comparing seedling survival to flowering success in cuttings (where selection at early life stages is bypassed), we show that genotypes with high seedling survival at the native elevation also tended to survive well at the novel elevation, but were genetically uncoupled from flowering success at the native elevation (genetic correlation near zero). This indicates that the persistence of genetic variation underlying adaptive potential does not require strong antagonistic trade-offs across native and novel environments, but can arise due to conditional neutrality where selection against genotypes that are adaptive in novel environments is not strong enough to purge them in native environments (Anderson et al. 2013). By showing that these genotypes are partially recessive, we provide evidence that being masked in heterozygotes reduces the potential for selection to remove them. If adaptive potential relies on partially recessive alleles as our results suggest, they may persist longer as standing variation. However, their increase in frequency in novel environments will likely be slower than additive or dominant alleles because their fitness advantages will only be partially expressed while rare (given most copies will occur in heterozygotes), limiting their ability to contribute to adaptation (Haldane 1927; Orr and Betancourt 2001). Whether these same mechanisms (weak selection and dominance) also contribute to the substantial standing genetic variation in fitness documented within native environments generally (Hendry et al. 2018; Bonnet et al. 2022) remains an open question.

Fluctuating selection provides another mechanism for maintaining genetic variation in the population: seasonal selection at the native elevation favoured different genotypes for seedling survival (Autumn versus Spring r_g_ = -0.49), consistent with balancing selection across temporally variable environments (Gillespie and Turelli 1989; Bürger and Gimelfarb 2002). However, we found no clear evidence that Autumn seedling survival predicted performance at the novel elevation, suggesting that the environmental axis driving seasonal selection at the native site did not align with the axis separating native from novel in this experiment. Fluctuating selection can therefore help preserve genetic (allelic) variance for seedling survival in native environments, without that variance necessarily corresponding to the particular genotypes needed for adapting to the novel high elevation environment (Connallon and Czuppon 2026). It is important to note that fluctuating selection could be created by factors other than temperature across seasons because many other variables change across seasons in different ways to changes across elevation, such as day length, precipitation or biotic communities. Such co-varying environmental factors could explain why seedling survival in Autumn did not predict performance at the novel elevation. Future work is therefore needed to test how environmental variation in the native range maintains genetic variation for adapting to novel environments.

Several alternatives that we could not test might also explain how genetic variation for increased adaptive potential is maintained in the population. Firstly, genotypes with higher fitness in novel environments could be held at mutation-selection balance and replenished as frequently as stabilising selection in native environments removes them (Lande 1975; Turelli 1984; Sharp and Agrawal 2018). However, several of our results are inconsistent with mutation-selection balance. For example, adaptive potential at the novel elevation was associated with genotype-specific fitness responses expressed consistently across experiments, life-history stages and generations, rather than the weak and inconsistent variance expected for random mutations with unpredictable fitness effects under mutation-selection balance (Houle 1992; Barton and Keightley 2002). Moreover, although mutation-selection balance is expected to enrich standing variation for partially recessive alleles, such alleles should be primarily deleterious (Sharp and Agrawal 2018), whereas the alleles underlying high fitness in the novel environment were conditionally beneficial. Instead, these patterns are more consistent with context-dependent fitness effects that reduce the efficiency of selection in the native environment. Secondly, it is also possible that spatial variation in selection among sites across the native range could favour different alleles, which could have contributed alleles that increased adaptive potential in the novel environment. However, we previously showed that genotypes with higher fitness in the novel environment were derived from several of our five sampling sites, without any evidence of local adaptation (Walter et al. 2023), making this possibility unlikely.

### Adaptive potential relies on rare genetic variants with context-dependent effects

Our results suggest that genotypes that increase adaptive potential may arise from partially recessive mutations that segregate in natural populations but whose positive fitness effects only emerge in novel environments. Adaptive capacity in novel environments is therefore likely to depend on genetic variants that are cryptic in native environments, making it difficult to predict responses to environmental change from performance under historical or native conditions alone (Hermisson and Wagner 2004; Walter et al. 2023). Given that we focus on a population exposed to a single novel environment, understanding the frequency and effect size of genotypes that increase adaptive potential across more environments, populations and species would help to predict the adaptive capacity of populations facing novel environments. For example, if adaptive potential depends on rare, recessive alleles of moderate to large effect, then adaptive capacity may be driven by a small number of variants (Hine et al. 2022).

An important implication of our results is that adaptive potential may rely on genetic variants whose fitness effects are rare, masked, or strongly context-dependent. Although balancing selection can maintain such variants under native conditions, their contribution to persistence during rapid environmental change will depend on their frequency, genetic architecture, and the population size. Even if balancing selection helps to maintain recessive alleles important for adapting to novel environments, populations may still lose these adaptive alleles through genetic drift during demographic decline before novel environments are encountered (Carlson et al. 2014; Orr and Unckless 2014). These issues are common in the context of evolutionary rescue, where persistence often depends on standing genetic variation that may be rare prior to environmental change (Gomulkiewicz and Holt 1995; Carlson et al. 2014; Bell 2017). Understanding how much adaptive capacity relies on rare and cryptic genetic variants, and how resilient these genetic variants are to demographic stochasticity, will be critical for predicting population persistence under rapid environmental change. While we focused on short-term fitness components, future work needs to quantify lifetime reproductive success across multiple generations to understand the potential for rare adaptive genotypes to colonise, persist and then maintain a stable population in a novel environment. This remains an ongoing challenge for field studies that test the potential for evolutionary rescue under environmental change (Carlson et al. 2014; Bell 2017).

## Conclusions

We provide field evidence that multiple processes can maintain genetic variation underlying adaptive potential in natural populations. We show that alleles important for persistence and potentially adaptation to a novel elevation segregate as partially recessive alleles, which are not strongly disadvantaged at later life-history stages in the native range, while changes in selection across seasons maintain genetic variation in fitness more broadly. Together, these findings suggest that adaptive potential can be maintained more broadly by variation in selection created by environmental heterogeneity, and more specifically because selection fails to remove genotypes whose effects on fitness are partially recessive or weakly coupled across life stages and environments. As a result, populations may harbour cryptic genetic variation that increases adaptive potential to environmental change, but that is not predictable from their performance in native conditions. Understanding the amount of such cryptic adaptive variation, and how reliably it contributes to adaptation, will be critical for predicting evolutionary responses to rapid environmental change.

## Materials and methods

### Breeding design

In 2017, we collected 72 individuals of *S. chrysanthemifolius* from five proximate sites (<5km) at elevations between 526-790masl around the base of Mount Etna. See Walter, *et al*. (Walter et al. 2023; Walter et al. 2024; Walter et al. 2026b) for a detailed description of the design. Briefly, we designated each individual as a sire or dam, and then mated 36 sires to 36 dams in blocks of 3×3, which produced 104 full-sibling families that we used in the experiments outlined below.

### Experiment 1: 2018 transplant of the breeding design as cuttings

Walter et al. (2023) describes the transplant in detail. We grew three individuals of each family in the glasshouse (n=312 individuals) until they were large plants, at which point we removed all their branches and cut them into smaller 4-5cm segments that we dipped in root hormone and allowed to establish as cuttings. We transplanted the cuttings at three elevations, a 500m home site in the native range, a 1500m site at the edge of the range in an apple and pear orchard, and a novel 2000m elevation on an old lava flow surrounded by pine trees. At each elevation, we transplanted ∼7-10 cuttings per glasshouse individual (n=c.2700 cuttings per elevation; N=8149 cuttings total) into 4-5 experimental blocks depending on the layout of the site. We transplanted the cuttings in Spring, allowed them to establish and grow for four months, and then measured fitness as the total number of flowers produced by each plant. This trait is routinely used to quantify fitness in short-lived perennials (Gross et al. 2004; Pujol et al. 2014), and has been linked to seed output in this system (Walter et al. 2023).

### Experiments 2-3: 2019 planting of seeds of the breeding design in Spring and Autumn

Walter, *et al*. (Walter et al. 2024; Walter et al. 2026b) describe the seed experiments in detail. Briefly, for each family of the breeding design, we glued 100 seeds each to a toothpick using non-drip super glue. In Spring (2019, experiment 2), at each of four elevations (with an additional 1000m elevation within the native range), we randomised 25 seeds per family into five blocks (n=540 seeds/block, n=2700 seeds/site, total N=10800 seeds). To prepare each block, we removed all plants and debris, and placed a plastic grid with 4cm square cells on the ground. Within each grid cell, we pressed a single toothpick into the ground so that the seed sat 1-2mm below the soil surface. We covered the blocks with 50% shadecloth and watered them for 2-3 weeks until germination ceased. We then tracked seedling emergence and mortality regularly until survival stabilised. To test whether different seasons favoured different genotypes, we repeated the seed experiment at 500m in the Autumn (2019, experiment 3) using the same replication of families, and the same experimental protocol. Seedling survival at 500m declined more rapidly in the Autumn compared to the Spring experiment (**Fig. S4**). Because survival is a binary trait, genetic variance in survival can only be estimated when survival is intermediate, we analysed survival when genetic variance could be estimated and was highest: 28 days post-sowing in Autumn and 49 days post-sowing in Spring at 500m, and 82 days post-sowing at 2000m. This criterion uses the genetic variance within each experiment to obtain the most robust estimate of genetic correlations across experiments. Direction of the genetic correlations were consistent on the survival days around those chosen.

### Statistical analyses for the first two hypotheses (weak and fluctuating selection)

All analyses were conducted in R version 4.5.2 (R Core Team 2025). To test whether weak and fluctuating selection help to maintain genetic variation underlying adaptive potential, we used generalised linear mixed models to connect genetic variation in fitness across life-history traits, elevations and seasons. We used *MCMCglmm* (Hadfield 2010) to apply

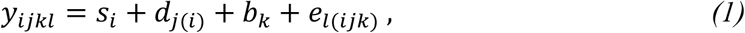

where we included the fitness in each environment as a multivariate response variable, where seedling survival was modelled as a threshold variable with a probit-link function, and flowering success as a Poisson variable with a log-link function. Random effects included sire (*s_i_*), dam nested within sire (*d*_j(*i*)_) and environmental block (*b_k_*), with *e_l_*_(*i*j*k*)_ the residual. For each application of equation 1, we specified unstructured covariance matrices for the sire component, which estimated additive genetic variances and covariances across environments. For the dam and block components, we only estimated variances in each environment, but not the covariances across environments. This was to reduce the number of parameters estimated, and because blocks and individuals were not measured in different environments, and so covariance estimation is not possible. To calculate additive genetic correlations across environments and life-history traits, for each application of equation 1 we extracted the posterior distribution for the sire component and used the *cov2cor* function to convert the covariance matrix into a correlation matrix.

We applied *MCMCglmm* models using a burn-in of 300000 iterations, a thinning interval of 1500 iterations and we saved 2000 MCMC samples as the posterior distribution for each parameter estimated. We used non-informative parameter-expanded priors, and checked that effective sample sizes were adequate, and that autocorrelation between samples did not exceed 0.1. To test the weak selection hypothesis, we included four response variables: flower production at 500m and 2000m (experiment 1, 2018), and seedling survival at 500m and 2000m (experiment 2, Spring 2019). This produced a 4×4 matrix that includes the covariances between the four performance estimates. To quantify statistical support, we calculated the probability of direction (pd) (Makowski et al. 2019), which is the proportion of the posterior distribution that has the same sign as the posterior mean. We report pd throughout, which ranges from 0.5 (no support for either direction) to 1. We describe estimates as credible when pd > 0.90. Because pd is one-sided and our 90% HPD intervals are two-sided, an estimate with pd between 0.90 and 0.95 can have a 90% HPD interval that overlaps zero. We therefore report both, so that the uncertainty in each estimate can be assessed directly. We also quantified differences in the correlations by conducting pairwise tests on all correlations in the matrix at each MCMC draw. To test the fluctuating selection hypothesis, we used the same approach but included seedling survival at 500m in Spring and Autumn (experiment 3, 2019) to estimate the genetic covariance between seasons. To test whether Autumn seedling survival predicted fitness in the novel environment, we then repeated the analysis and included Autumn seedling survival, and then performance at 2000m as seedlings (Spring 2019) and as flower production (2018).

### Experiment 4: 2020 transplant of cuttings derived from crossing contrasting genotypes

To understand the genetic architecture of AP genotypes, and to test the extent to which AP genotypes are heritable, we crossed within and between AP and HR genotypes and transplanted cuttings from offspring grown in the glasshouse. As described in Walter et al. (2026a), we identified the genotypes (individuals from the breeding design) that showed the biggest difference in their fitness response to elevation in experiment 1 (2018 transplant) – see **Fig. 3**. In the original study, we found a negative genetic correlation in fitness of -0.11 between the native 500m site and the 2000m novel elevation at the population level. To select the genotypes with contrasting fitness responses to elevation based on this negative correlation, we chose 12 Adaptive Potential (AP) genotypes that showed the highest relative fitness at 2000m, which we contrasted with 13 Home Range (HR) genotypes that showed the highest relative fitness at the native elevation.

We conducted crosses to produce 20 AP×AP families, and 16 HR×HR families. Given this species is hermaphroditic, individuals can produce reciprocal crosses. To ensure each side of the cross between AP and HR genotypes was independent, we used different genotypes to produce 17 AP×HR families, and 23 HR×AP families. To conduct crosses, we placed perforated bread bags over the plants to prevent pollinator contamination. We then rubbed mature flowerheads from the pollen donor (sire) on the flowerhead of the pollen acceptor (dam). Fertilised flowerheads were then labelled, covered with jewellery bags, and the seeds collected once mature.

We grew one individual from each family in the glasshouse using the same standard growth protocol as in Walter et al. (2023). Once plants were fully grown, we took 32 cuttings per genotype and transplanted 8 cuttings at each transplant site, randomised into 4 blocks (76 genotypes, 8 cuttings/elevation, n=608 cuttings/elevation; total N=2432 cuttings). Cuttings were transplanted 19^th^ June (2020) using the same protocol described in experiment 1. After five months (November), we collected all flowerheads from each plant in separate paper bags and counted them as our proxy for fitness. We then collected two leaves from each plant, which we weighed, and then scanned and measured using the program *Lamina* (Bylesjo et al. 2008). We used four traits to represent leaf morphology and investment: leaf area, complexity 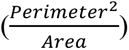, number of leaf indents divided by perimeter, and Specific Leaf Area 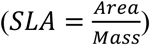.

### Statistical analyses for the recessive alleles hypothesis

To test whether AP genotypes were recessive, we compared the fitness and phenotype of AP×AP and HR×HR genotypes to AP×HR genotypes using *MCMCglmm* to apply the generalised linear mixed model

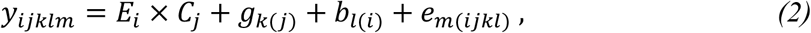

where the interaction between the fixed effects of transplant elevation (*E_i_*) and cross type (*C*_j_; AP×AP, AP×HR or HR×HR) tests whether the different crosses vary in their response to changes in elevation. Random effects included the *k*th genotype within cross type (*g_k_*_(j)_), and the *l*th block within transplant elevation (*b_l_*_(*i*)_). For the genotype random effect, we estimated variation in each elevation, as well as the covariance across elevation, resulting in a 4×4 matrix. We modelled fitness (number of flowers) with a Poisson distribution (log-link function), and we log-transformed the leaf traits to account for variance decreasing with mean at higher elevations. We applied the models with the same priors as equation 1, except that we used a more constrained parameter-expanded prior for the block variances (αV = 1 rather than 50), because there were few blocks per elevation (four in experiment 4) from which to estimate the block variance. We also reduced the total iterations to 550000. From each model, we extracted the posterior distributions of the fixed effects to test for credible elevation × cross-type interactions, indicating that differences between AP×AP and HR×HR crosses changed between native and novel elevations.

To test whether AP genotypes are recessive, we calculated the potence ratio 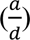, where *a* represents additivity (half the parental difference) and *d* the deviation due to dominance (hybrid deviation from the midparent) (Falconer 1960; de Vienne and Fievet 2020). We calculated the potence ratio relative to the HR×HR parent, which means that zero represented additivity, and a value of 1 represents full dominance towards HR genotypes. Calculating the potence ratio for each MCMC draw provided uncertainty in the estimate.

We estimated plasticity for each genotype from the output of equation 2. First, we extracted the posterior distribution for each genotype measured at each elevation. For each MCMC draw, we added the appropriate fixed effects, and then calculated plasticity using

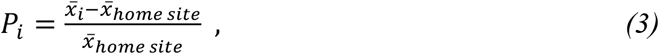

where plasticity (*P_i_*) is estimated for each genotype as the difference in mean phenotype between the 500m native site and the *i*th elevation, standardised by the native site mean (Valladares et al. 2006; Walter et al. 2026a). This captures plasticity as the elevational change in magnitude and direction (negative values reflect a trait decrease) of the phenotype relative to the native site (Anderson et al. 2021). We then summarised plasticity for each genotype class (AP×AP, AP×HR and HR×HR) as the mean across the appropriate genotypes, which provided the posterior distribution of each genotype class at each elevation.

## Acknowledgements

We are very grateful to everyone who helped with fieldwork in the original experiments, particularly Enrico la Spina, Giuseppe Pepe and Maria Majorana, as well as Alessandro Barbato, Octavia Brayley, Guy Burstein, Maria Castrogiovanni, Stefania Catara, Carmen Impelluso, Jessica Menzies, Morgan Millen and Daniel Ward. We are also grateful to Piante Faro (Giarre, Italy) for providing us with glasshouse facilities that made this study possible, and we thank Giuseppe Riggio for generously providing us access to the 1000m field site. We thank Jia Huan Liew and Luke Yates for providing comments on early versions.

## Author contributions

GW designed the experiments with JB, and input from SC, SH, JC and AC. GW, DT, GE and AC conducted the experiments. GW analysed the data, and then wrote the manuscript with JB, and input from all other authors.

## Supplementary material

**Fig. S1.**
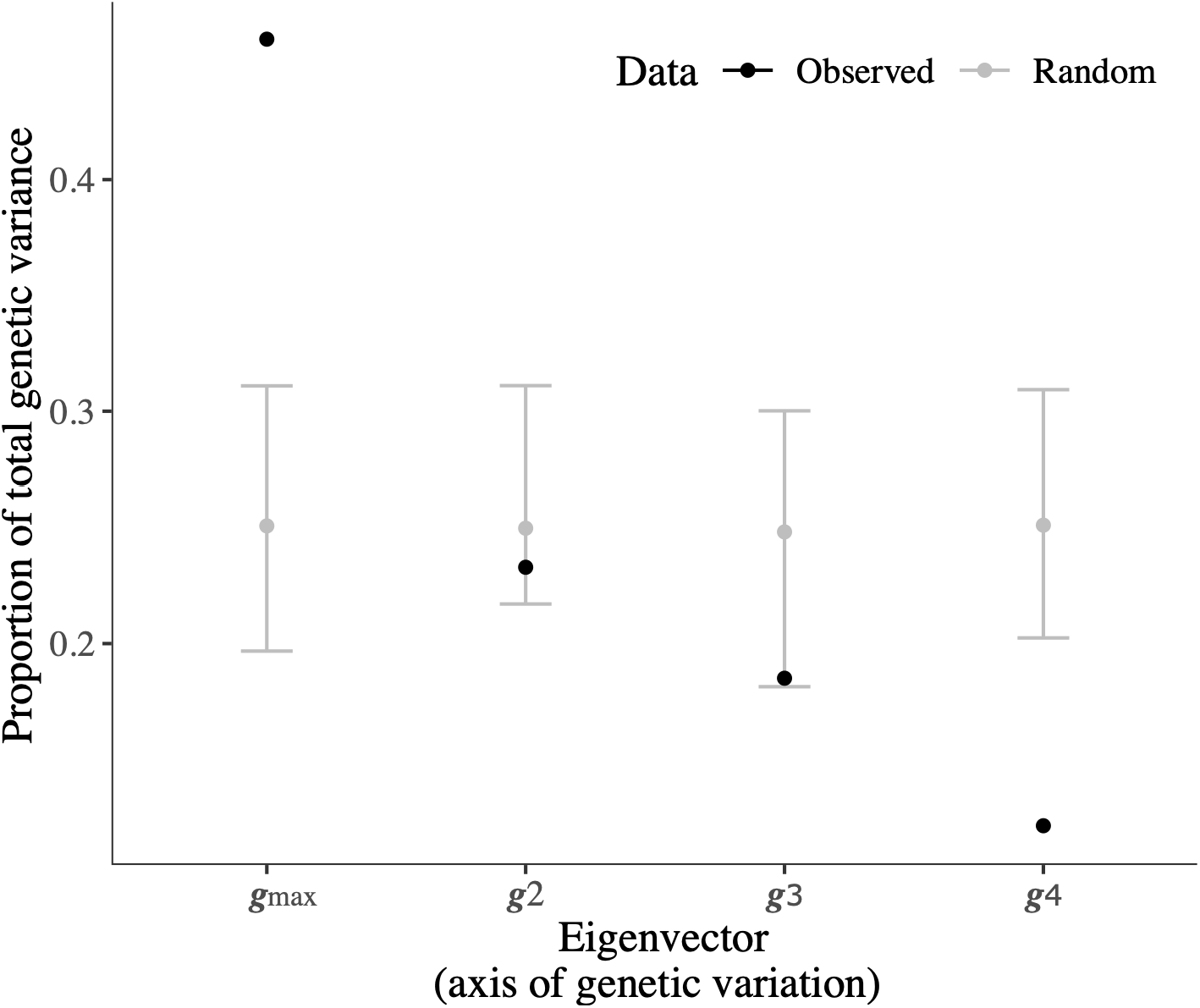
We used a permutation test to test whether axes of additive genetic variation underlying the four field performance traits (seedling survival and flowering success in both native and novel elevations) described more genetic variation than expected under a suitable null distribution. We created the null distribution by randomising the pedigree across the fitness (survival or flowering success) data within experimental blocks (i.e., for each block within each elevation), and then re-applying the model to each randomised dataset. The grey points and error bars represent the 95% confidence interval of the null distribution, calculated across the mean of each model applied to a single randomisation. Where the mean of the observed model (black points) exceeds the null distribution provides evidence that the genetic variation captured by that particular axis is greater than expected at random. We found strong statistical support for the leading axis, ***g***_max_, but no other axis.

**Table S1.**
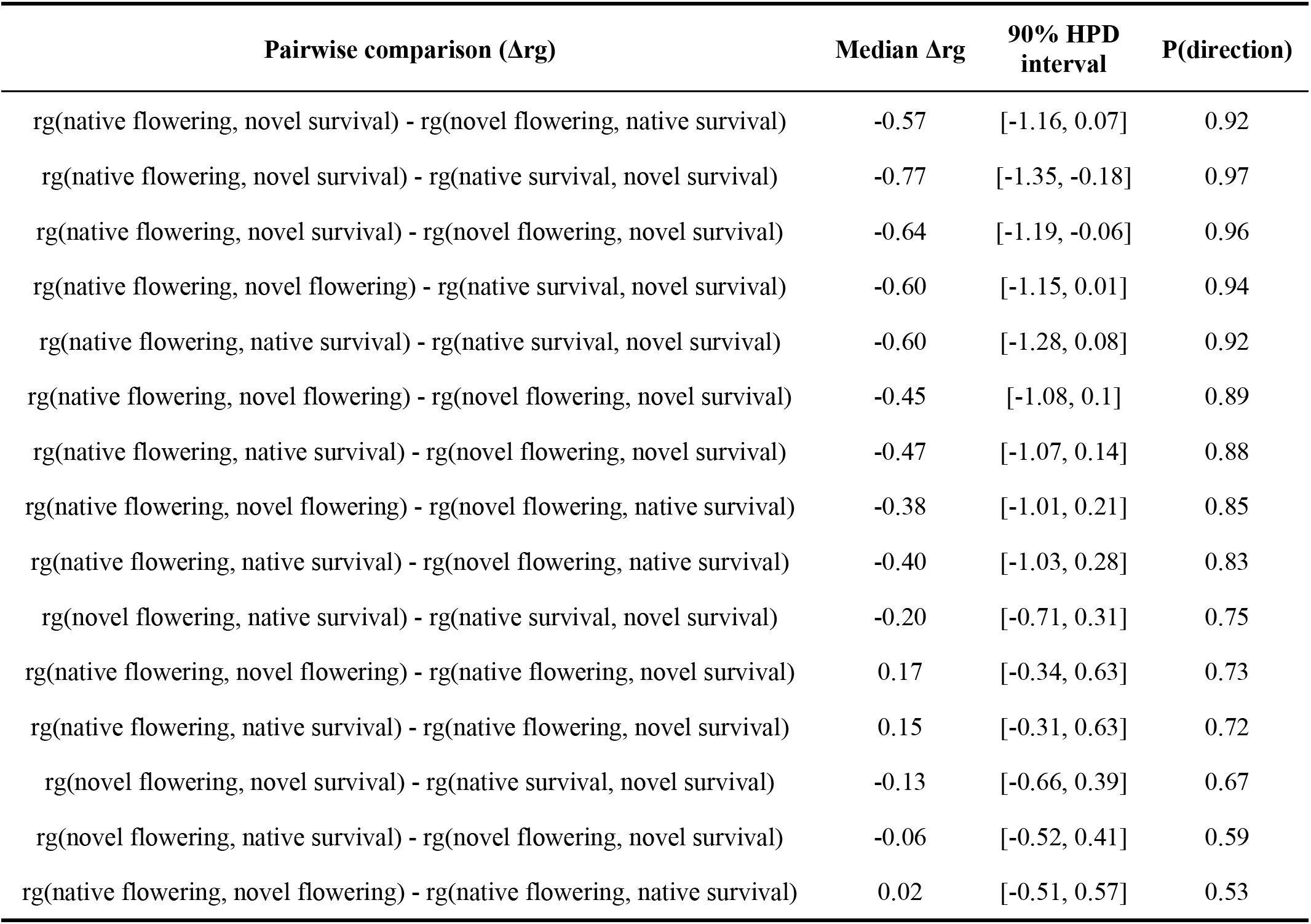
Pairwise comparisons of genetic correlations in **Table 1A**.

| Pairwise comparison (Δrg) | Median Δrg | 90% HPD interval | P(direction) |
| --- | --- | --- | --- |
| rg(native flowering, novel survival) - rg(novel flowering, native survival) | -0.57 | [-1.16, 0.07] | 0.92 |
| rg(native flowering, novel survival) - rg(native survival, novel survival) | -0.77 | [-1.35, -0.18] | 0.97 |
| rg(native flowering, novel survival) - rg(novel flowering, novel survival) | -0.64 | [-1.19, -0.06] | 0.96 |
| rg(native flowering, novel flowering) - rg(native survival, novel survival) | -0.60 | [-1.15, 0.01] | 0.94 |
| rg(native flowering, native survival) - rg(native survival, novel survival) | -0.60 | [-1.28, 0.08] | 0.92 |
| rg(native flowering, novel flowering) - rg(novel flowering, novel survival) | -0.45 | [-1.08, 0.1] | 0.89 |
| rg(native flowering, native survival) - rg(novel flowering, novel survival) | -0.47 | [-1.07, 0.14] | 0.88 |
| rg(native flowering, novel flowering) - rg(novel flowering, native survival) | -0.38 | [-1.01, 0.21] | 0.85 |
| rg(native flowering, native survival) - rg(novel flowering, native survival) | -0.40 | [-1.03, 0.28] | 0.83 |
| rg(novel flowering, native survival) - rg(native survival, novel survival) | -0.20 | [-0.71, 0.31] | 0.75 |
| rg(native flowering, novel flowering) - rg(native flowering, novel survival) | 0.17 | [-0.34, 0.63] | 0.73 |
| rg(native flowering, native survival) - rg(native flowering, novel survival) | 0.15 | [-0.31, 0.63] | 0.72 |
| rg(novel flowering, novel survival) - rg(native survival, novel survival) | -0.13 | [-0.66, 0.39] | 0.67 |
| rg(novel flowering, native survival) - rg(novel flowering, novel survival) | -0.06 | [-0.52, 0.41] | 0.59 |
| rg(native flowering, novel flowering) - rg(native flowering, native survival) | 0.02 | [-0.51, 0.57] | 0.53 |

**Fig. S2.**
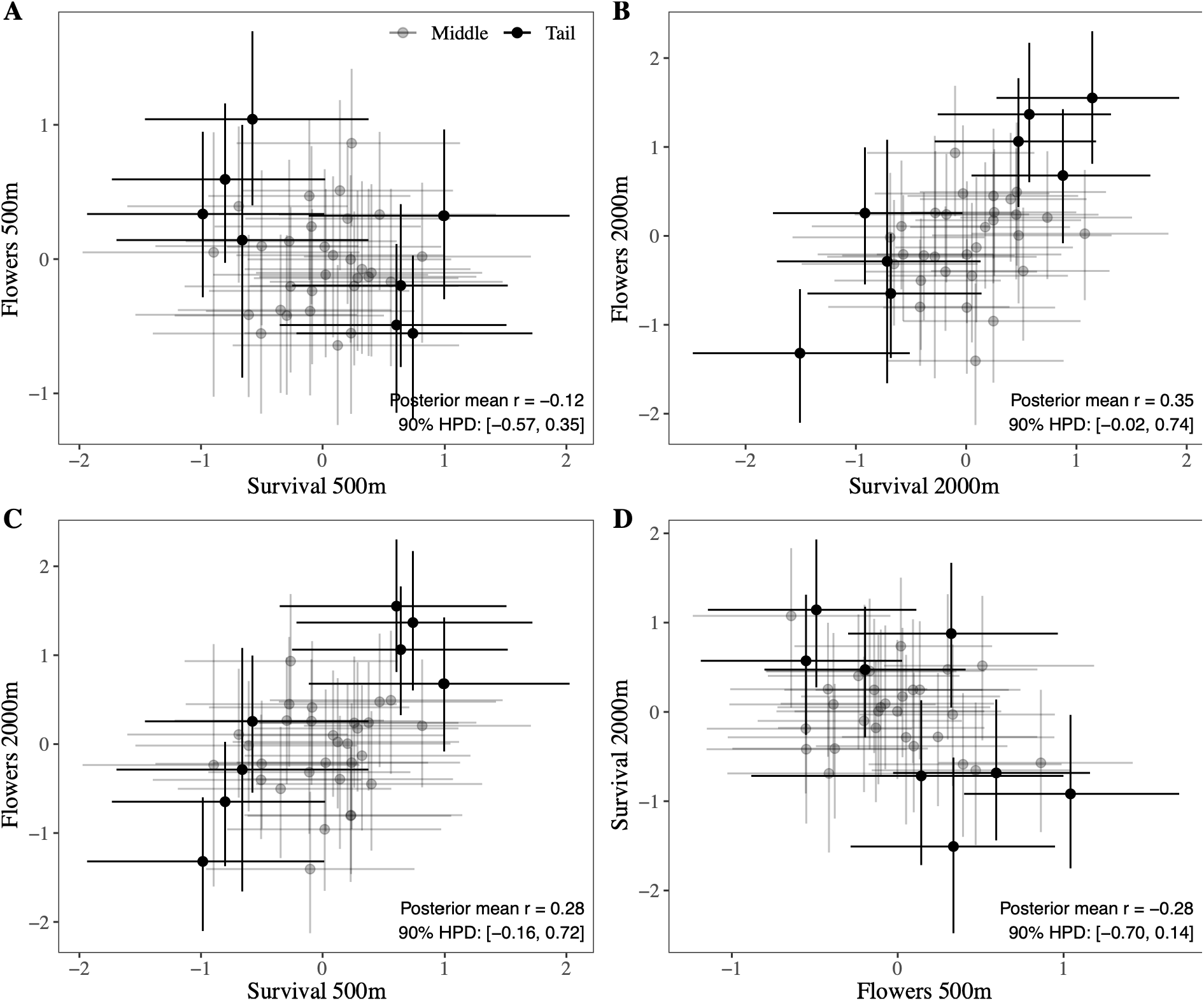
Sire BLUPs underlying the genetic correlations in **Table 1** and **Fig. 1**, but including 90% HPD credible intervals for each BLUP. The sires from the tails of the multivariate fitness distribution are emphasised in black, and the remaining genotypes in grey. Although the posterior distribution of the genetic correlations overlap zero, there are still differences between the tails of the distribution suggesting that genotypes with high and low fitness in panels **C-D** show credible differences.

**Fig. S3.**
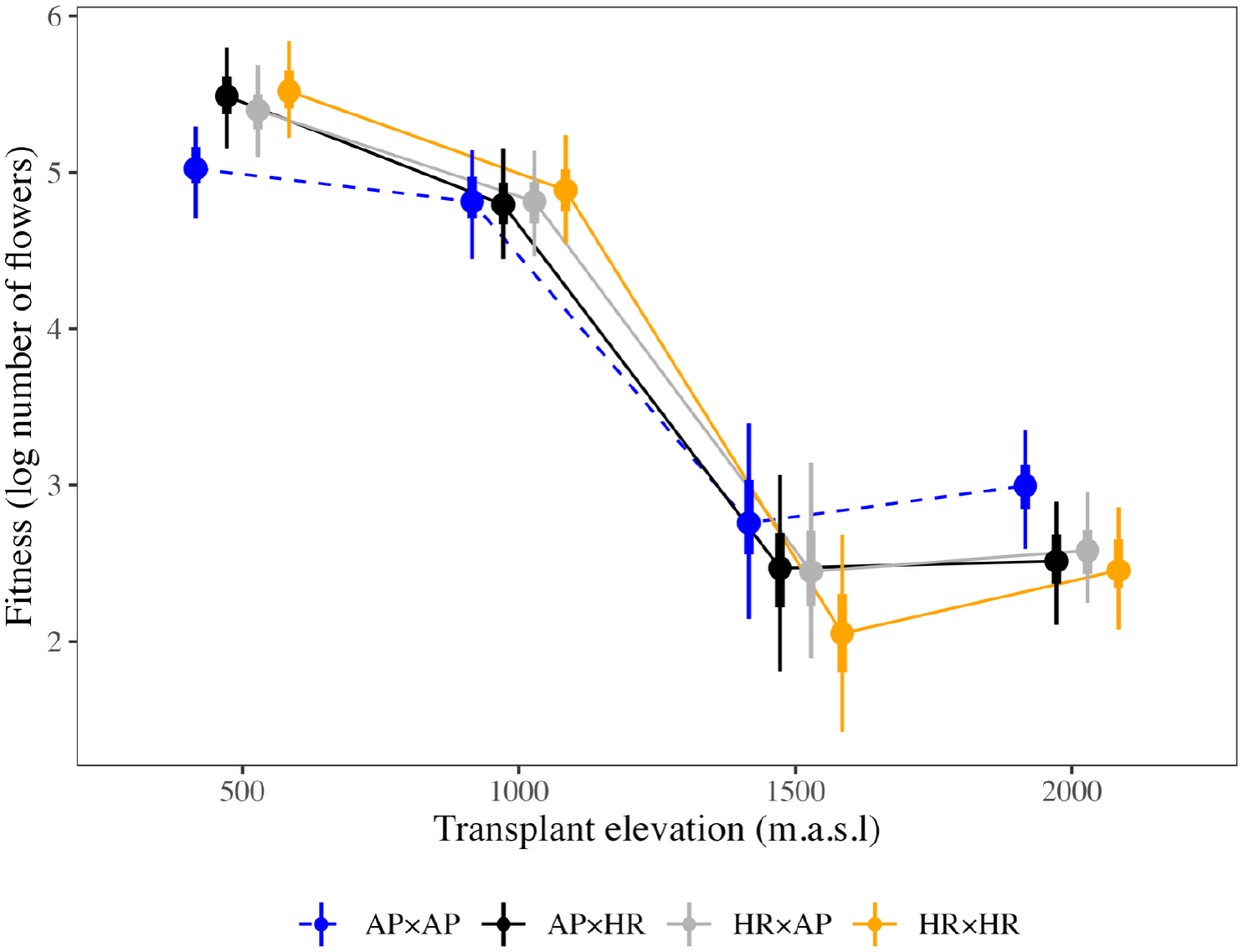
Fitness of cuttings as the log number of flowers for AP×AP and HR×HR genotypes as per **Fig. 3**. Also included are the reciprocal hybrids between them: AP×HR (black) and HR×AP (grey) genotypes, which show similar fitness at all elevations, suggesting the direction of the cross does not change fitness. Points represent the posterior median and error bars the 50% (thick) and 90% (thin) HPD credible intervals.

**Fig. S4.**
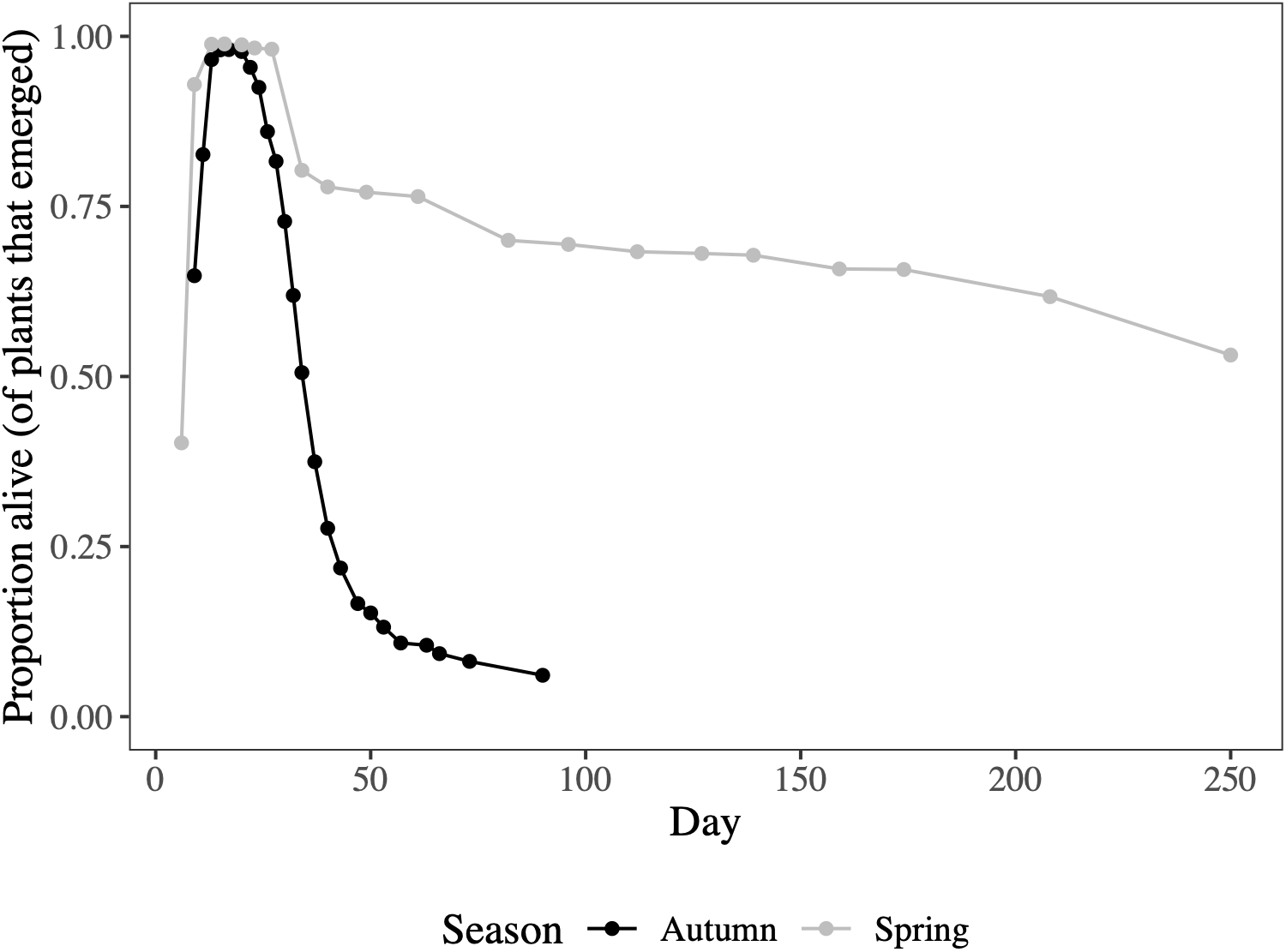
Comparing survival for the Spring (grey) and Autumn (black) seedling experiments.

**Table S2.** Summary table for the test of pairwise interactions between genotype and elevations for fitness. Interactions are calculated as whether the difference between AP×AP and HR×HR change across elevations. P is the probability of direction (pd), i.e., P(<0) or P(>0).

| Year | Contrast | Mean (90% HPD) | P |
| --- | --- | --- | --- |
| 2018 | (AP - HR 500m) - (AP - HR 1500m) | -0.77 (-1.06, -0.49) | 1.00 |
|  | (AP - HR 500m) - (AP - HR 2000m) | -2.33 (-2.71, -1.93) | 1.00 |
|  | (AP - HR 1500m) - (AP - HR 2000m) | -1.56 (-1.92, -1.24) | 1.00 |
| 2020 | (AP×AP - HR×HR 500m) - (AP×AP - HR×HR 1000m) | -0.43 (-0.74, -0.09) | 0.99 |
|  | (AP×AP - HR×HR 500m) - (AP×AP - HR×HR 1500m) | -1.20 (-1.65, -0.78) | 1.00 |
|  | (AP×AP - HR×HR 500m) - (AP×AP - HR×HR 2000m) | -1.04 (-1.50, -0.59) | 1.00 |
|  | (AP×AP - HR×HR 1000m) - (AP×AP - HR×HR 1500m) | -0.78 (-1.15, -0.43) | 1.00 |
|  | (AP×AP - HR×HR 1000m) - (AP×AP - HR×HR 2000m) | -0.61 (-0.97, -0.28) | 1.00 |
|  | (AP×AP - HR×HR 1500m) - (AP×AP - HR×HR 2000m) | 0.16 (-0.27, 0.56) | 0.74 |

**Table S3.** Summary table for the test of pairwise interactions between genotype and elevations for the leaf traits. Interactions are calculated as whether the difference between AP×AP and HR×HR change across elevations. P is the probability of direction (pd), i.e., P(<0) or P(>0).

| Trait | Contrast | Mean (90% HPD) | P |
| --- | --- | --- | --- |
| Leaf area | (AP×AP - HR×HR 500m) - (AP×AP - HR×HR 1000m) | -0.13 (-0.31, 0.06) | 0.87 |
|  | (AP×AP - HR×HR 500m) - (AP×AP - HR×HR 1500m) | -0.37 (-0.59, -0.17) | 1.00 |
|  | (AP×AP - HR×HR 500m) - (AP×AP - HR×HR 2000m) | -0.25 (-0.44, -0.05) | 0.98 |
|  | (AP×AP - HR×HR 1000m) - (AP×AP - HR×HR 1500m) | -0.24 (-0.44, -0.06) | 0.98 |
|  | (AP×AP - HR×HR 1000m) - (AP×AP - HR×HR 2000m) | -0.12 (-0.27, 0.06) | 0.88 |
|  | (AP×AP - HR×HR 1500m) - (AP×AP - HR×HR 2000m) | 0.12 (-0.07, 0.32) | 0.85 |
| Leaf complexity | (AP×AP - HR×HR 500m) - (AP×AP - HR×HR 1000m) | -0.04 (-0.13, 0.05) | 0.79 |
|  | (AP×AP - HR×HR 500m) - (AP×AP - HR×HR 1500m) | -0.18 (-0.29, -0.07) | 1.00 |
|  | (AP×AP - HR×HR 500m) - (AP×AP - HR×HR 2000m) | -0.10 (-0.18, 0.00) | 0.95 |
|  | (AP×AP - HR×HR 1000m) - (AP×AP - HR×HR 1500m) | -0.14 (-0.24, -0.02) | 0.98 |
|  | (AP×AP - HR×HR 1000m) - (AP×AP - HR×HR 2000m) | -0.05 (-0.16, 0.05) | 0.80 |
|  | (AP×AP - HR×HR 1500m) - (AP×AP - HR×HR 2000m) | 0.08 (-0.03, 0.19) | 0.88 |
| Indentation | (AP×AP - HR×HR 500m) - (AP×AP - HR×HR 1000m) | 0.02 (-0.05, 0.09) | 0.69 |
|  | (AP×AP - HR×HR 500m) - (AP×AP - HR×HR 1500m) | 0.04 (-0.05, 0.13) | 0.73 |
|  | (AP×AP - HR×HR 500m) - (AP×AP - HR×HR 2000m) | 0.04 (-0.03, 0.12) | 0.81 |
|  | (AP×AP - HR×HR 1000m) - (AP×AP - HR×HR 1500m) | 0.01 (-0.09, 0.11) | 0.60 |
|  | (AP×AP - HR×HR 1000m) - (AP×AP - HR×HR 2000m) | 0.02 (-0.06, 0.10) | 0.65 |
|  | (AP×AP - HR×HR 1500m) - (AP×AP - HR×HR 2000m) | 0.00 (-0.09, 0.09) | 0.54 |
| SLA | (AP×AP - HR×HR 500m) - (AP×AP - HR×HR 1000m) | 0.00 (-0.08, 0.06) | 0.50 |
|  | (AP×AP - HR×HR 500m) - (AP×AP - HR×HR 1500m) | 0.06 (-0.02, 0.14) | 0.87 |
|  | (AP×AP - HR×HR 500m) - (AP×AP - HR×HR 2000m) | 0.05 (0.00, 0.12) | 0.94 |
|  | (AP×AP - HR×HR 1000m) - (AP×AP - HR×HR 1500m) | 0.06 (-0.03, 0.13) | 0.86 |
|  | (AP×AP - HR×HR 1000m) - (AP×AP - HR×HR 2000m) | 0.05 (-0.01, 0.12) | 0.92 |
|  | (AP×AP - HR×HR 1500m) - (AP×AP - HR×HR 2000m) | 0.00 (-0.08, 0.08) | 0.51 |

**Fig. S5.**
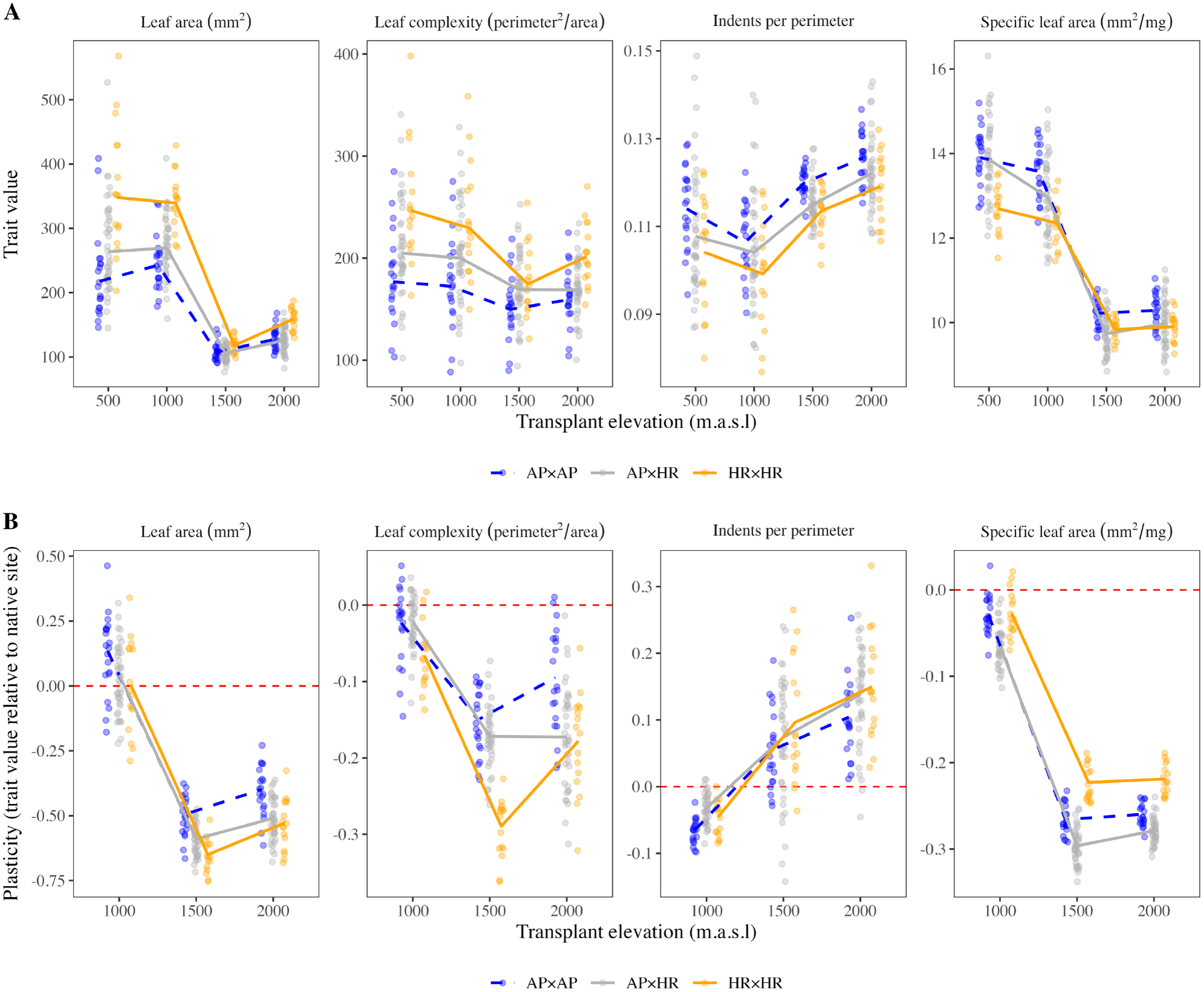
Genotype estimates for trait values and plasticity across elevations. Small points show the posterior median for each genotype for (A) trait values and (B) plasticity relative to the native 500m site, coloured by cross type. Lines connect the posterior medians for each cross type, as presented in Fig. 4.

